# Color Polymorphism is a Driver of Diversification in the Lizard Family Lacertidae

**DOI:** 10.1101/2020.08.27.270207

**Authors:** Kinsey M. Brock, Emily Jane McTavish, Danielle L. Edwards

## Abstract

Color polymorphism – two or more heritable color phenotypes maintained within a single breeding population – is an extreme type of intra-specific diversity widespread across the tree of life but rarely studied in a comparative framework. Color polymorphism is thought to be an engine for speciation, where morph loss or divergence between distinct color morphs within a species results in the rapid evolution of new lineages, and thus, color polymorphic lineages are expected to display elevated diversification rates. Lizards of the family Lacertidae have evolved multiple lineages with color polymorphism, but lack of a complete and robust phylogeny for the group has made comparative analysis difficult. Here, we produce a comprehensive species-level phylogeny of the lizard family Lacertidae to reconstruct the evolutionary history of color polymorphism and test if color polymorphism has been a driver of diversification. Accounting for phylogenetic uncertainty, we estimate an ancient macroevolutionary origin of color polymorphism within the Lacertini tribe (subfamily Lacertinae). Color polymorphism most likely evolved several times in the Lacertidae and has been lost at a much faster rate than gained. Evolutionary transitions to color polymorphism are associated with shifts in increased net diversification rate in this family of lizards. Taken together, our empirical results support long-standing theoretical expectations that color polymorphism is a driver of diversification.

---

Understanding how diversity is generated and maintained within and across species is a fundamental goal of evolutionary biology. Color polymorphism, or the presence of two or more distinct genetically determined color phenotypes within a breeding population (Huxley 1955; Gray & McKinnon 2007), exemplifies extraordinary intra-specific phenotypic diversity. Despite its usefulness in developing population genetic theory (Ford 1945; Huxley 1955; Svensson 2017), relatively little is known about the evolutionary origins and macroevolutionary consequences of color polymorphism (Gray & McKinnon 2007; Jamie & Meier 2020). A longstanding hypothesis is that the presence of different genetically-based color morphs within populations may promote speciation (West-Eberhard 1986; Jonsson 2001; Gray & McKinnon 2007; Corl et al. 2010a,b; Hugall & Stuart-Fox 2012; McLean et al. 2014). In this scenario, a group of distinct morphs is established within a population, phenotypic variation among morphs gradually accumulates, and over time isolated populations with different morphs diverge, facilitating speciation (West-Eberhard 1986). Empirical studies spanning fish (Seehausen & Schluter 2004; Allender et al. 2003), birds (Hugall & Stuart-Fox 2012), and reptiles (Corl et al. 2012) all suggest the evolution of multiple variable morphs as an impetus for speciation. After more than 50 years of research, we have amassed a body of literature comprising theoretical studies, reviews, and empirical work on individual species that suggest color polymorphism should be an engine of speciation (Jonsson 2001; Gray & McKinnon 2007; Forsman et al. 2008; McLean et al. 2014). However, few comparative studies exist that test this claim (but see Hugall & Stuart-Fox 2012).

Theoretical and empirical work have generated hypotheses that predict the evolutionary history and persistence of color morphs and a positive relationship between color polymorphism diversification rates that can be tested with comparative methods. Evolutionary gains of novel genetically-based color morphs are expected to occur at a slower rate than they are lost (Corl et al. 2010b; Hugall & Stuart-Fox 2012). Empirical studies of color polymorphic species have consistently found that geographic variation in the number and types of morphs is usually explained by morph loss from populations, with little evidence for gaining morphs back (Corl et al. 2010b). Forsman et al. (2008) suggest that color morphs with alternative co-adaptations could promote the utilization of broader environmental resources and range expansion, less vulnerability to environmental change, robustness to extinction and more evolutionary potential for speciation via divergent natural selection. Alternatively, if color morphs are maintained by negative frequency dependent selection (Sinervo & Lively 1996), where color morph fitness and frequencies may oscillate dramatically, rare morphs may be more easily lost from populations, and the process of divergence may progress quickly toward speciation. Thus, we expect that color polymorphism is more easily lost than gained, that alternative color morphs provide standing phenotypic variation for evolutionary processes to promote speciation, and that these speciation events likely result in color polymorphic ancestors giving rise to monomorphic lineages. However, not all color polymorphisms evolved or are maintained by the same mechanisms (Gray & McKinnon 2007), which adds complexity for determining the relationship between color polymorphism and rates of speciation and extinction. Idiosyncrasies in the mechanisms generating and maintaining color polymorphism matter a great deal when trying to make predictions about how microevolutionary processes generate macroevolutionary patterns. What the literature lacks and needs are comparative studies of large species groups to test long-standing hypotheses about the evolution of color polymorphism - the rate at which it evolves relative to how quickly it is lost, the macroevolutionary trait patterns it generates, and its relationship, or not, to diversification.

Phylogenetic comparative methods use reconstructions of the historical relationships among taxa (i.e., phylogenies) to compare and contrast features of organisms given shared ancestry, and have been a powerful tool for the study of evolution (Felsenstein 1985). Comparative methods such as ancestral state reconstruction (Revell 2013) and diversification models (Rabosky et al. 2014; O’Meara & Beaulieu 2016) are commonly used to model the evolutionary history of traits and their influence on macroevolutionary dynamics (Morlon 2014). Given color morph dynamics observed from empirical studies of populations (Sinervo & Lively 1996; Corl et al. 2010b; Runemark et al. 2010), we expect that the evolution of color polymorphism is rare, and the rate of loss of color polymorphism exceeds the rate of gain across the phylogeny. Trait evolution and its relationship to diversification can be jointly assessed with time-calibrated phylogenies and state-dependent speciation and extinction models (Maddison et al. 2007; O’Meara & Beaulieu 2016). State-dependent speciation and extinction models are a birth-death process that deal with binary traits, and test whether binary trait transitions happen at speciation events (nodes in a phylogeny) or along branches. If color polymorphic species are engines of speciation, then net diversification rates (speciation – extinction) should be greater within color polymorphic lineages compared to monomorphic lineages (Hugall & Stuart-Fox 2012). Accurately estimating or distinguishing between alternative speciation and extinction scenarios on phylogenies with rate-varying diversification models is difficult (Louca & Pennell 2020), and strong arguments have been made against trying to identify one true diversification history given uncertainty. However, we can construct suites of alternative diversification models that incorporate trait data and encode biological knowledge in parameter specifications to tease out what effects speciation and extinction have on model estimates (Maddison et al. 2007; Beaulieu & O’Meara 2016). Further, we can leverage Bayesian methods to construct a random sample of phylogenies from our posterior distribution of possible trees (i.e., evolutionary histories) and use this sample to compare diversification hypotheses and parameter estimates under different evolutionary scenarios and asses model adequacy.

The family Lacertidae (Oppel 1811) is an excellent group for a comparative study on the evolution of color polymorphism as the group is relatively speciose and multiple species spanning several genera are known to be color polymorphic (Vercken & Clobert 2008; Huyghe et al. 2009a,b; Runemark et al. 2010; Brock et al. 2020 in press). The Lacertidae is the most speciose family of squamates in the Western Palearctic comprising around 320 species distributed across Africa, Asia, and Europe (Arnold et al. 2007). Biologists have identified a similar throat color polymorphism in several genera across the family Lacertidae (Fig. 1), including *Iberolacerta monticola* (López et al. 2009), *Podarcis* species (*P. gaigeae*, Runemark et al. 2010; *P. melisellensis*, Huyghe et al. 2009a,b; *P. muralis*, Pérez i de Lanuza et al. 2019), and *Zootoca vivipara* (Vercken & Clobert 2008). Though color polymorphism has been studied in individual species, we lack a broader understanding of how this trait has evolved at the macroevolutionary scale in any group. Progress on understanding of the evolution of this trait through comparative explorations across the Lacertidae have been hampered by a lack of comprehensive and robust phylogenies (Fu 2000; Arnold et al. 2007). Historically the mitochondrial genome has been used to infer evolutionary relationships at the generic-level in lacertids (Fu 2000), with more recent studies using a combination of both mitochondrial and nuclear genes (Pyron et al. 2013; Baeckens et al. 2015; Garcia-Porta et al. 2019). Missing genetic data from under-sampled geographic regions in combination with gene tree discordance has been a significant historical hurdle for inferring family-wide species-level phylogenies (Fu 2000; Arnold et al. 2007). However, the opportunity to leverage new data and analytical approaches mean the Lacertidae is now an ideal group for a comparative study on the evolution of color polymorphism.

**Figure 1.**
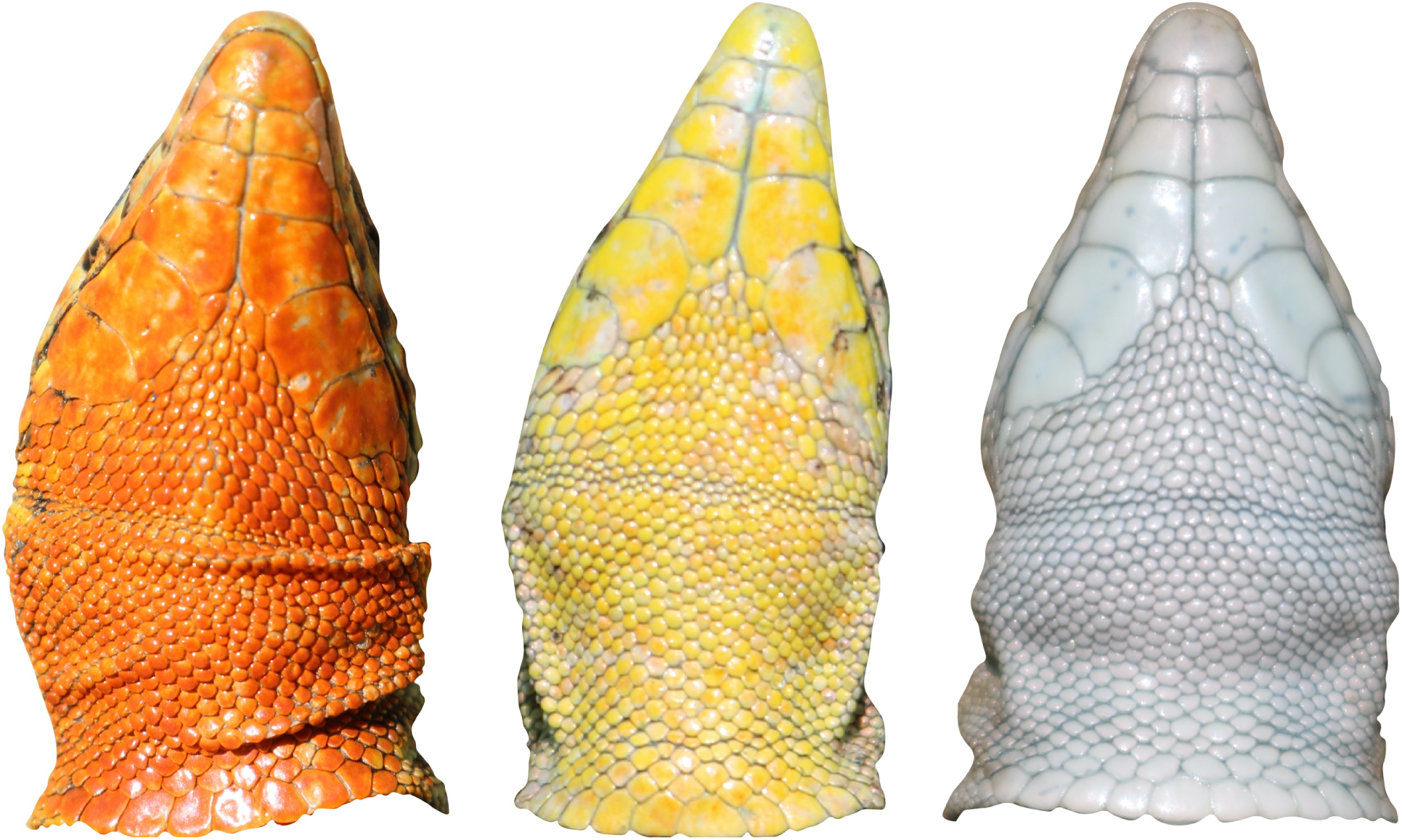
Throat color polymorphism in *Podarcis erhardii*, a Mediterranean lacertid species. In lacertid lizards, species spanning several genera exhibit a similar color polymorphism. Across all lacertid species in our study, color polymorphism is expressed at adulthood as colorful badges ventrally on the throat or belly region.

In this study, we use the evolutionary relationships between species in the family Lacertidae to understand the evolutionary history of color polymorphism and test whether the evolution of this trait impacts diversification rate shifts. We build a comprehensive species-level phylogeny of the Lacertidae, including 262 species from all 42 described genera. We apply a multispecies coalescent approach that accounts for potential individual gene histories among species (in contrast to approaches used in previous studies, Fu 2000; Arnold et al. 2007; Pavlicev & Mayer 2009). Using this species-level tree, we conduct the first family-wide investigation of color polymorphism and assess long-standing hypotheses concerning the evolutionary consequences of this trait. Specifically, we address the following questions: (1) What is the evolutionary history of color polymorphism in the Lacertidae? Color polymorphism is rare, and even within species populations seem to lose color morphs more easily than they gain them (Corl et al. 2010b). Given this, we expect that color polymorphism evolved once deep in the tree and was subsequently lost several times. Thus, we also expect that the evolutionary rate of losing color polymorphism far exceeds the rate at which it is gained. (2) Do color polymorphic lineages have elevated rates of diversification compared to non-color polymorphic lineages? Mathematical models suggest that the evolution of multiple color forms within a population may increase evolutionary potential for speciation through increased utilization of diverse habitats and resources, and greater ability to successfully colonize and expand their range relative to monomorphic populations (Forsman et al. 2008). Therefore, we predict that evolutionary transitions to color polymorphism are associated with elevated diversification rates.

## MATERIALS & METHODS

### Phylogenetic Inference of Family Lacertidae

#### GenBank data

We used publicly available sequence data from GenBank to build our tree (Clark et al. 2016). We first pulled all Lacertidae single gene sequences from GenBank and identified five genetic markers for which there were at least 100 species with data for that marker. This filtering resulted in three mitochondrial genes (12s: N = 178 species; 16s: N = 189 species; cytochrome b: 239 species) and two nuclear genes (mos proto-onco gene: N = 142 species; recombination activating gene (RAG) 1: N = 117 species). Genes selected from the mitochondrial genome have been used extensively for phylogenetic reconstruction of Lacertidae in the past (Fu 2000; Edwards et al. 2012; Pyron et al. 2013; Baeckens et al. 2015; Baeckens et al. 2017), and the RAG-1 nuclear gene has been shown to evolve at a greater rate than other commonly used nuclear markers, which may allow for greater confidence both deep within the tree and at the tips (Portik et al. 2012; Edwards et al. 2012). To ensure taxonomic validity, sequences were georeferenced to verify they fell within species distributions as described on the regularly updated lacertid database, AG Lacertiden (www.lacerta.de, maintained by The Arbeitsgemeinschaft Lacertiden within the Deutsche Gesellschaft für Herpetologie und Terrarienkunde). Overall, we retrieved gene sequences for 262 lacertid species – approximately 82% of described lacertid diversity according to the AG Lacertiden database, covering all currently described genera (Arnold et al. 2007). Details of genetic data used per species and GenBank accession numbers for each genetic marker are provided in Supplemental 1. Sequences were aligned separately in AliView (v.1.25) using MUSCLE (v.3.8.425). Gene sequence alignments were then assessed for appropriate models of molecular evolution for tree inference using the Bayesian Information Criterion in jModelTest (v.2.1.10) (Darriba et al. 2012).

#### Species tree inference and divergence dating estimation

We employed a multi-locus coalescent approach for species tree inference as individual gene histories can vary within closely related species (Maddison 1997; McCormack et al. 2009). To do this with our five locus alignments, we used the full Bayesian method of species tree estimation in BEAST2 using the *BEAST template (v.2.5.1) (Heled & Drummond 2010; Bouckaert et al. 2014). We unlinked all site and molecular clock models, and linked the gene tree models for mtDNA alignments only. We used the GTR+Γ+I site model for each gene according to model comparison results from jModelTest with all frequencies set to empirical and no additional parameters estimated for computational efficiency. For all genes, we specified an uncorrelated relaxed molecular clock with a log-normal distribution that assumes each branch has its own independent rate (Drummond et al. 2006). The species tree population size function was set to linear with constant root and population mean set to 1.0. We set the species tree prior to a Yule Model (pure-birth) with a log-normal distribution on the species birth-rate. Prior distributions for all gene clock-rate priors were set to exponential, and the population mean prior was set to log-normal. To time-calibrate the phylogeny, we used the ‘Sampled Ancestors’ package in BEAUti2 to generate monophyletic taxon set hyper-priors according to fossil information from the literature (Hipsley et al. 2009). Time-calibrated outgroup nodes following Hipsley *et al*. (2009) include: (1) *Sphenodon punctatus* – *Cnemidophorus tigris*, 228.0 Mya (Sues & Olsen 1990), (2) *Cnemidophorus tigris* – *Rhineura floridana*, 113.0 Mya (Nydam & Cifelli 2002), and (3) *Rhineura floridana* – *Gallotia galloti*, 64.2 Mya (Sullivan 1985). All fossil hyper-priors were offset by the above dates, given a log-normal distribution with a mean of 1.0 and standard deviation of 1.25, and constrained to be monophyletic. We ran six independent species tree analyses with the same data and XML configuration for 1 billion MCMC generations and stored every 30,000th sampled tree. Posterior distributions of trees for the six independent BEAST2 runs were combined in BEAST2’s logCombiner (v.2.5.1) with the first 20% of trees discarded as burn-in and resampled at a lower frequency for a total posterior density of 12,000 trees. A final maximum clade credibility tree, the tree from the reduced posterior sample that had the maximum sum of posterior probabilities on its n - 2 internal nodes, was generated in BEAST2’s TreeAnnotator for use in comparative analyses.

### The Evolution of Color Polymorphism and State-Dependent Diversification

#### Color polymorphism data

We scored the presence or absence of color polymorphism for all described extant lacertid species (N = 320), including those not represented in our phylogeny. Examination of all extant taxa, including species lacking available genetic data and not represented in our phylogeny, was necessary to account for trait estimation proportions downstream in our state-dependent speciation and extinction (SSE) models. We scored color polymorphism from several georeferenced sources to ensure taxonomic validity, including online databases with photographs (www.inaturalist.org, www.lacerta.de), scientific literature (Huyghe et al. 2009a,b; López et al. 2009; Runemark et al. 2010), and field guides (Valakos et al. 2008; Speybroeck et al. 2016). We *a priori* restricted our investigation to color polymorphism in the truest sense – that is, species with multiple coexisting color morphs in the same geographic location. Further, we focused on color polymorphism in the same trait on the same region of the body, as this is likely to share both a similar underlying genetic mechanism and subject to similar selective pressures across species (Andrade et al. 2019). In lacertid lizards, color polymorphism is expressed as colorful badges on the throat and colors vary from species to species (Runemark et al. 2010). Species were coded as color polymorphic if we could adequately identify they met all of the following criteria: 1) variation in color located on the throat, 2) variation in throat color is not result of ontogenetic color change and is present in adults, 3) individuals from the same location exhibit at least two different throat colors, and 4) variation in throat color is not strictly sexually dimorphic, males and females both exhibit at least two different color types. We were able to collect color polymorphism data for all described lacertids and thus had no missing trait data. Altogether, we identified 43 color polymorphic species spanning 10 genera.

#### Ancestral state reconstruction of color polymorphism in the Lacertidae

To understand the evolutionary history of color polymorphism in lacertids, we performed ancestral state reconstruction on our time-calibrated maximum clade credibility tree. We also ran the same set of analyses on a recently published lacertid tree inferred by Garcia-Porta et al. (2019) that differed somewhat in taxon representation and topology to test if our results were robust to phylogenetic uncertainty. Ancestral state reconstructions were carried out in R using a maximum likelihood approach with the ‘ace’ function in the ‘ape’ package (Paradis & Schliep 2018). We also estimated ancestral states and the associated uncertainty of our discretely valued color polymorphism trait at each node in the tree using a continuous-time Markov chain model, or Mk model in the ‘phytools’ package (Revell 2012). We fit a single-rate model with equal transition rates and an all-rates-different model that allows backward and forward transition rates between states to have different values and compared model log-likelihood scores using a likelihood ratio test with ? set to 0.05 *a priori* to select the best fit model. We summarized results from our final all-rates-different Mk model by mapping the empirical Bayesian posterior probability pie charts for each node (Fig. 2).

**Figure 2.**
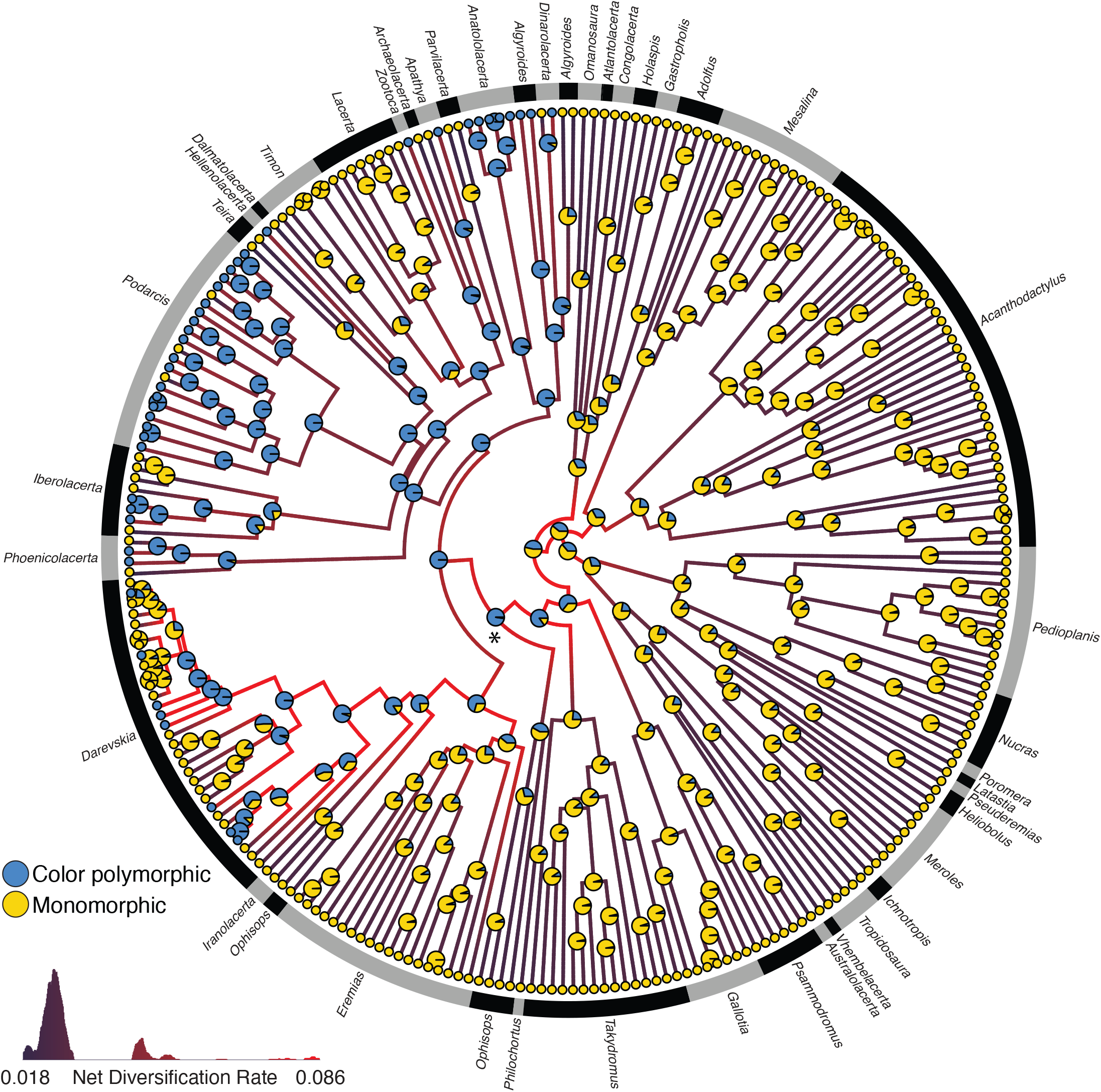
Evolutionary relationships of 262 species from the family Lacertidae. Phylogeny is the maximum clade credibility (MCC) tree calculated from the posterior of a full-Bayesian species tree inference from three mitochondrial and two nuclear loci. Genera are labelled and separated by black and gray bars. Observed color polymorphism character states for extant taxa are given in circles at the tips (blue indicates observed species is color polymorphic, yellow indicates observed species is monomorphic). Empirical Bayesian posterior probability pie charts at the nodes indicate inferred ancestral state probabilities. An asterisk denotes the oldest common ancestor with 99% probability of being color polymorphic. Branches of the phylogeny are painted with net diversification rates from the best fit HiSSE model, and indicate the estimated tempo of net diversification, ranging from slowest rate (0.018) to fastest rate (0.086).

#### State-dependent diversification models

To test our hypothesis that evolutionary transitions to color polymorphism are associated with elevated diversification rates, we used state-dependent speciation and extinction (SSE) models (Maddison et al. 2007). An advantage of SSE models is joint estimation of trait transitions and diversification rates (Maddison et al. 2007; Beaulieu & O’Meara 2016). The original binary state-dependent speciation and extinction (BiSSE) model calculates the probability that a group of extant species evolved as observed at the tips given a phylogenetic tree and a binary character under a simple model of evolution with six parameters (Maddison et al. 2007). The parameterization of a basic BiSSE model specifies two speciation rates (a rate for when a lineage is in state 0, and a rate for when a lineage is in state 1), two extinction rates (for lineages in state 0 and state 1), and two rates of character state transition (from state 0 to state 1 and *vice versa*). The hidden-state speciation and extinction (HiSSE) model framework is an extension of BiSSE that specifies additional parameters to account for diversification rate heterogeneity that is not associated with the observed trait (Beaulieu & O’Meara 2016). These “hidden states” represent unmeasured characters that could affect diversification rate estimates for the measured observed character (Beaulieu & O’Meara 2016). The SSE model framework is statistically advantageous because BiSSE models are nested within HiSSE models, and maximum likelihood inference can be used to estimate a suite of alternative models and their parameters for subsequent hypothesis tests (see below). Biologically, SSE models are desirable for our study because we are interested in both the evolutionary history of color polymorphism (historic transitions to and from color polymorphism) and if this character, or something unmeasured, is associated with increased speciation and extinction or not at all.

We performed BiSSE and HiSSE model tests in R with the ‘HiSSE’ package (Beaulieu & O’Meara 2016). We constructed a suite of character-dependent (BiSSE and HiSSE), character-independent models (CID-2 and CID-4), and null models to test alternative hypotheses related to the evolution of color polymorphism and diversification rates in the family Lacertidae (Table 1). Briefly, the CID-2 and CID-4 models are BiSSE and HiSSE character-independent models, respectively. The CID-2 and CID-4 models contain the same number of distinct turnover and extinction fraction parameters that can vary across the tree as their analogous BiSSE and HiSSE models, but CID-2 and CID-4 explicitly specify that diversification is not linked to the observed character state (Beaulieu & O’Meara 2016). The BiSSE and HiSSE null models also contain the same number of transition rates as the BiSSE and HiSSE models, but the null models specify a constant rate of diversification across the tree (number of distinct turnover rates = 1). For all six SSE models run on our time-calibrated species tree, we used the same estimated proportion of extant species (82% of all non-color polymorphic and 100% of color polymorphic extant species are represented in our tree) and did not constrain the root character state. Full SSE model parameterizations are given in Supplemental 2. Nested SSE models were compared using AICc scores, ΔAIC scores, and Akaike weights (Table 1) (Burnham & Anderson 2002).

**Table 1.** Summary of color polymorphism SSE hypotheses, model parameterization, and associated model fits. Models are listed in best-fit order according to AIC value. A HiSSE model, with state-dependent diversification and hidden states, best explained the data.

| Model | Log Likelihood | AIC | $\Delta$ AIC | W AIC | N Distinct Turnover Rates | N Distinct Extinction Fractions | N Distinct Transition Rates |
| --- | --- | --- | --- | --- | --- | --- | --- |
| HiSSE | -1238.173 | 2496.346 | 0 | 0.999 | 4 | 1 | 5 |
| BiSSE | -1249.974 | 2509.948 | 13.602 | 0.001 | 2 | 1 | 2 |
| HiSSE null | -1249.422 | 2512.843 | 16.497 | < 0.0001 | 1 | 1 | 5 |
| BiSSE null | -1260.179 | 2523.294 | 26.948 | < 0.0001 | 1 | 1 | 2 |
| HiSSE CID-4 | -1253.647 | 2528.357 | 32.011 | < 0.0001 | 4 | 1 | 3 |
| BiSSE CID-2 | -1260.179 | 2532.358 | 36.012 | < 0.0001 | 2 | 1 | 3 |

#### Trait simulations and SSE model adequacy

The main disadvantage of SSE models is the potential issue of choosing between model A and model B when model C is true (Caetano et al. 2018). To address this potential issue, we conducted a simulation study (Portik et al. 2019) to identify the rate at which a character-dependent model of diversification (BiSSE or HiSSE) is falsely chosen as the correct model out of all six SSE models with uncorrelated simulated color polymorphism trait data on our empirical phylogeny. To simulate color polymorphism character data on our species tree, we used the ‘phytools’ (v.0.6-60) ‘sim.history’ function (Revell 2012). For trait data simulation, we used the same all-rates-different Q transition matrix from our ancestral state reconstruction analysis and the inferred character state posterior probability from the root of all lacertids from our empirical data. We extracted the root state probabilities from our fitted model and used them to specify simulation root states (probability that the MRCA of all lacertids is color polymorphic = 64%, not color polymorphic = 36%). We then simulated character data on our empirical tree 10,000 times and randomly selected 1,000 simulations where at least 10% of tips were color polymorphic for SSE model adequacy investigation. We then ran the same five SSE models from our empirical study on our 1,000 simulations and extracted AIC scores for model comparison and adequacy evaluation (Fig. 3). Given that the simulated trait data are uncorrelated, we expect character-independent (CID-2 and CID-4) or null models (BiSSE null and HiSSE null) to have the lowest AIC scores most of the time.

**Figure 3.**
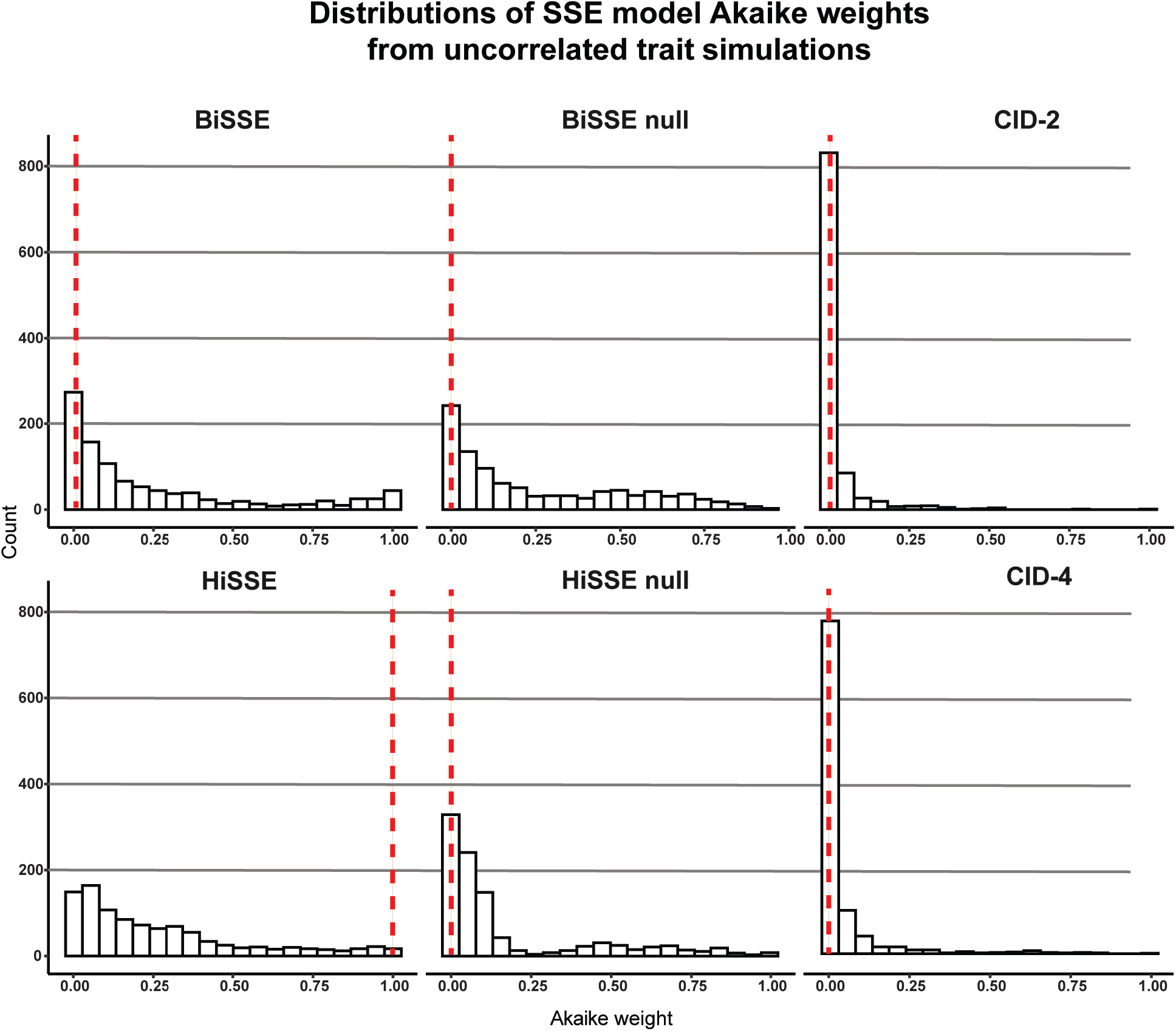
Distributions of Akaike model weights for six state-speciation extinction (SSE) models run on 1,000 simulated color polymorphism trait datasets. All simulations and models were performed with our empirical phylogeny. The horizontal axis (Akaike weight) refers to the relative probability of that SSE model compared to the other five competing SSE models run on the same uncorrelated simulated color polymorphism trait data. The dotted vertical lines indicate model weights from our empirical phylogeny and observed color polymorphism trait data for that same SSE model. Low model weight indicates relatively low support for the that hypothesis, and high model weight indicates greater support.

#### Phylogenetic uncertainty and SSE model comparison

A phylogenetic tree represents one hypothesis of evolutionary relatedness. Biological conclusions drawn from phylogenetic comparative methods are influenced by uncertainty in the timing and topology of those relationships, with the potential for misleading conclusions based on mis-estimating the true diversification history (Louca & Pennell 2020). To understand the robustness of our conclusions based on phylogenetic comparative analyses carried out on our maximum clade credibility tree, we performed a sensitivity analysis using 1,000 randomly selected trees from the posterior distribution of 12,000 trees from our Bayesian tree inference. Here, we ran the same six SSE models (Table 1) with the same parameterizations on 1,000 possible phylogenies with our observed color polymorphism trait data. We extracted AIC scores for SSE models run on each tree for model comparison (Fig. 4). Finally, we extracted model parameter estimates from a subset of phylogenies from the posterior sample of 1,000 trees to understand how uncertainty in phylogeny affects net diversification and extinction fraction estimates (N = 50 phylogenies, Table 2).

**Table 2.**
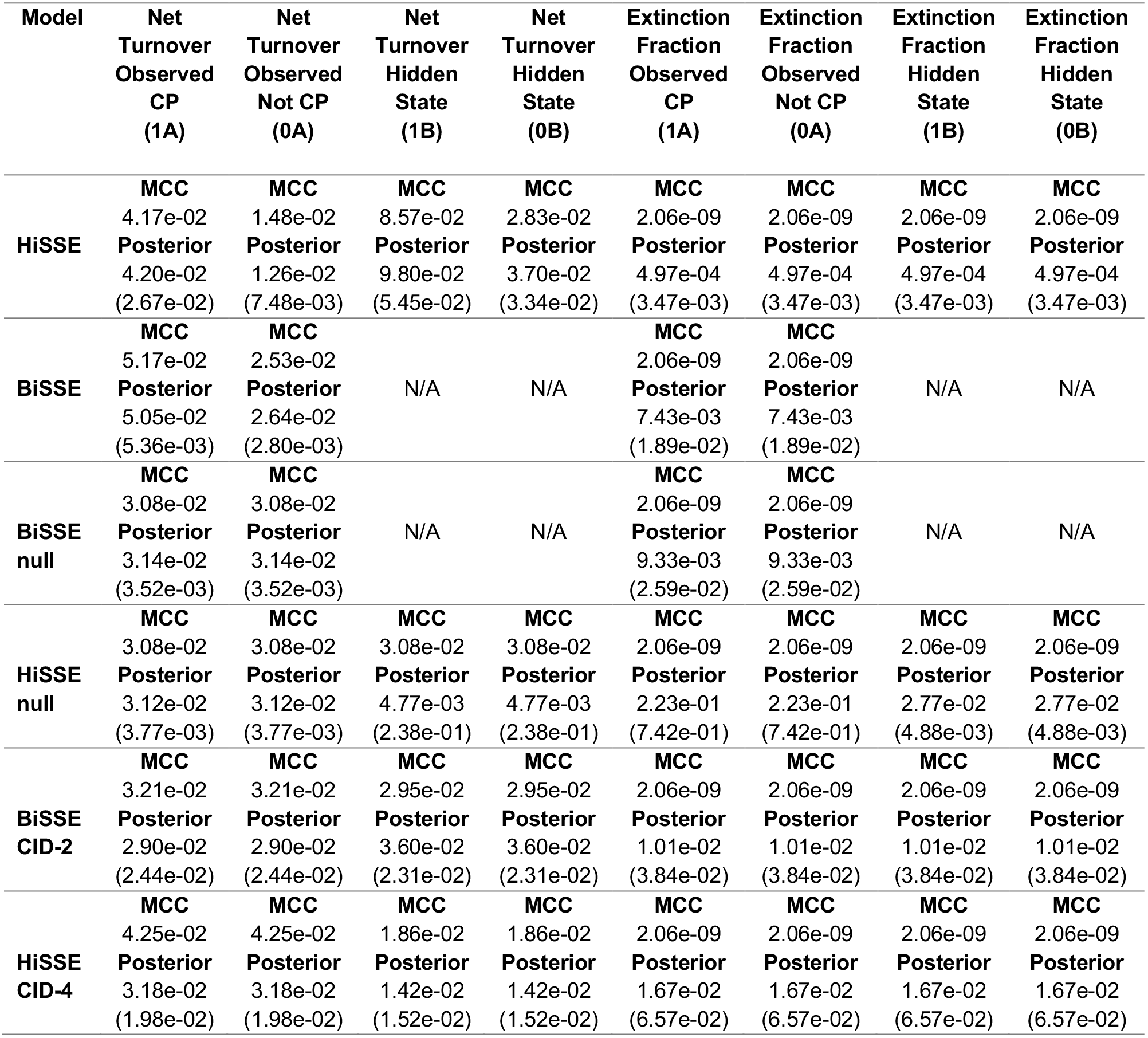
SSE model parameter output from maximum clade credibility species tree and poster sample of 50 trees. Net turnover and extinction fraction parameter estimates from the maximum clade credibility (MCC) species tree (Figure 2) are given for each SSE model under MCC. SSE model parameter estimates from 50 phylogenies extracted from the posterior are given under Posterior where the first value is the average and the value in parentheses is the standard deviation of the estimate. For all six SSE models, turnover parameter estimates from the MCC tree are contained within one standard deviation from the mean posterior estimate.

**Figure 4.**
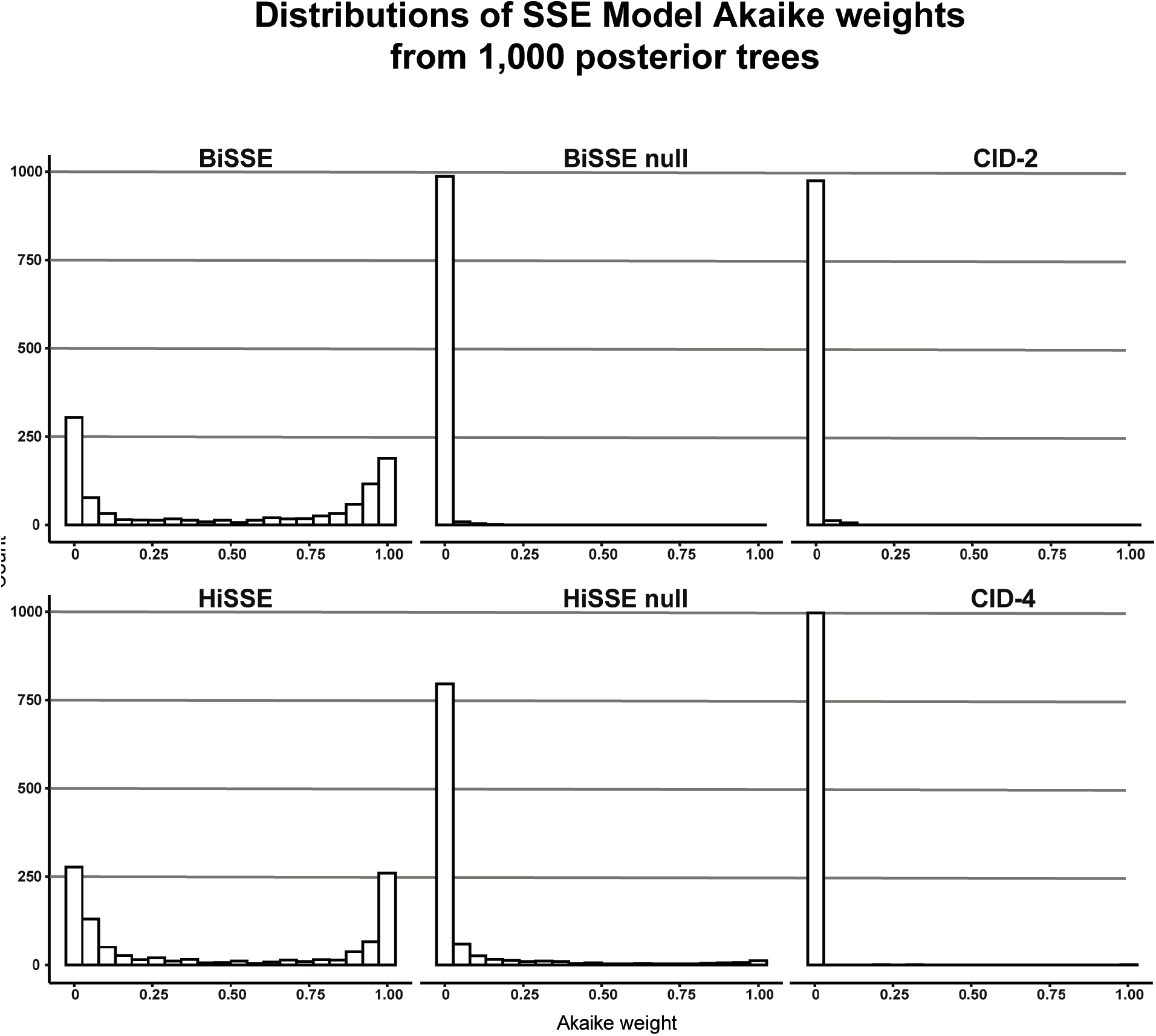
Distributions of Akaike model weights for six SSE models run on 1,000 phylogenetic trees randomly selected from posterior distribution of possible trees and our empirical color polymorphism trait dataset. A higher model weight indicates greater relative support for that state-speciation extinction hypothesis.

All data and code for this project are available on Dryad.

## RESULTS

### Phylogeny of the Lacertidae

Phylogenetic inference from the combined six independent MCMC runs with 20% burn-in converged well (ESS values for posterior, likelihood, species coalescent, and all five gene trees > 200). Evolutionary relationships presented in our maximum clade credibility tree (Fig. 2, all node posterior probabilities given in Supplemental 3), recovered from our multi-locus full Bayesian species tree inference are generally consistent with other family-wide phylogenies (Hipsley et al. 2009; Garcia-Porta et al. 2019). The subfamily Gallotiinae, comprising *Gallotia* and *Psammodromus* genera, grouped together monophyletically consistent with previous studies (Arnold et al. 2007; Garcia-Porta et al. 2019). However, our inference placed Gallotiinae nested within Lacertinae with low node support (posterior probability = 0.2377), which follows results from Fu (2000), but is in contrast to the hypothesis that the Gallotiinae and Lacertinae are two separate monophyletic subfamilies that comprise the Lacertidae (Arnold et al. 2007; Garcia-Porta et al. 2019). The two tribes within the subfamily Lacertinae (as most recently reviewed with 620bp of mtDNA and 64 morphological characters from 59 nominal species by Arnold et al. 2007), the Lacertini and Eremiadini, were not reciprocally monophyletic. The Lacertini tribe (Oppel 1811; Arnold et al. 2007) that usually comprise 18 genera from Europe, northwest Africa, and southwest and east Asia largely grouped together, with 15 of 18 genera forming a monophyletic clade containing *Algyroides, Anatololacerta, Apathya, Archaeolacerta, Dalmatolacerta, Dinarolacerta, Hellenolacerta, Iberolacerta, Lacerta, Parvilacerta, Phoenicolacerta, Podarcis, Teira, Timon*, and *Zootoca*. The *Darevskia* and *Iranolacerta* genera that belong to the Lacertini tribe grouped monophyletically sister to the aforementioned Lacertini, but also include the *Eremias*, which are traditionally placed in the Eremiadini tribe. The last Lacertini genus, the *Takydromus*, which contains 24 species that have a far east distribution spanning eastern China, Japan, and southeast Asia grouped monophyletically but separate from the other Lacertini. The Eremiadini tribe diverge from other lacertids deep in our phylogeny, with the speciose *Acanthodactylus* and *Meroles* genera forming a monophyletic clade, the Afrotropical genera *Pedioplanis, Nucras, Poromera, Latastia, Pseuderemias, Heliobolus, Meroles, Ichnotropis, Vhembelacerta*, and *Australolacerta* forming a monophyletic clade. The remaining Eremiadini genera from Equatorial Africa, *Adolfus, Gastropholis, Holaspis, Congolacerta, Atlantolacerta*, and *Omanosaura* form a separate clade similar to Arnold et al. (2007). Most genera formed monophyletic groups with the exception of *Algyroides, Eremias*, and *Ophisops*.

### The evolutionary history of color polymorphism in the Lacertidae

We identified 43 color polymorphic extant lacertid species spanning 10 out of 42 currently described genera. The 10 genera containing color polymorphic species are all within the sub-family Lacertinae: *Algyroides* (2 of 4 spp.), *Anatololacerta* (5 of 5 spp.), *Apathya* (1 of 2 spp.), *Darevskia* (8 of 30 spp.) *Dinarolacerta* (1 of 2 spp.), *Hellenolacerta* (1 of 1 sp.), *Iberolacerta* (3 of 8 spp.), *Phoenolacerta* (2 of 4 spp.), *Podarcis* (19 of 23 spp.), and Zootoca (1 of 1 sp.). All 43 color polymorphic lacertid species are represented in our phylogeny and comparative analyses. We ran comparative analyses on both our inferred species and the tree proposed by Garcia-Porta et al. (2019) which show consistent results, so here we present only the results from our phylogeny and provide detailed results from the Garcia-Porta et al. (2019) tree in Supplemental 4.

A likelihood ratio test supported the more heavily parameterized all-rates-different transition probability matrix where all possible transitions between states receive distinct parameters best fit the data (ARD log-likelihood = −78.86843, ER log-likelihood = − 90.37925, likelihood ratio test: p < 0.001, df = 1).

Ancestral state reconstruction using the all-rates-different transition matrix revealed that the most recent common ancestor of all lacertids was most likely not color polymorphic (Scaled likelihood at the root, color polymorphic = 64%, not color polymorphic = 0.36%, Fig. 2). Results from ancestral state reconstruction also suggest that color polymorphism is an ancient trait within lacertids, and is highly likely to have evolved by the most recent common ancestor of *Ophisops* and *Dinarolacerta* (Scaled likelihood common ancestor was color polymorphic = 99%, Fig. 2, denoted with an asterisk), but could have evolved even earlier in the common ancestor of *Takydromus* and *Dinarolacerta*.

Results from sampling 1,000 character histories conditioned on the all-rates-different transition matrix detected 38 total character changes in color polymorphism, and suggest that color polymorphism most likely evolved seven times in this family of lizards, and was subsequently lost 31 times throughout the evolutionary history of the Lacertidae (Fig. 2). Estimates of evolutionary transition rates from mono-to polymorphism (Rate index estimate = 0.0009, std. error = 0.0010 - 0.0004) were much lower than rates of transition from poly-to monomorphism (Rate index estimate = 0.02118, std. error = 0.0212 - 0.0038), suggesting color polymorphism is more easily lost than gained.

### State-dependent diversification in the Lacertidae

Results from SSE analyses on the maximum clade credibility tree support a character state-dependent diversification model with hidden states, or HiSSE model (Akaike weight = 0.999, Fig. 2, Table 1). For the best fit model, turnover parameter estimates were asymmetrical between color polymorphic and non-color polymorphic lineages (MCC parameter estimates, Table 2). Estimated net diversification rates were higher in observed color polymorphic lineages (HiSSE observed 1A color polymorphic net diversification rate = 0.0417, observed 0A monomorphic net diversification rate = 0.0148, Table 2). Parameter estimates for character transitions from color polymorphism to monomorphism were much higher than transitions from monomorphism to color polymorphism, providing further evidence that color polymorphism is more easily lost than gained (HiSSE observed 1A color polymorphic transition rate to monomorphic 0A = 0.0166, observed monomorphic 0A to color polymorphic 1A transition rate = < 0.0001). The other state-dependent diversification model without hidden states, BiSSE, had the second best fit model (BiSSE ΔAIC = 13.602, Akaike weight = 0.001, Table 1). Parameter estimates from the BiSSE model also detected higher rates of net diversification at evolutionary transitions to color polymorphism (BiSSE observed 1A color polymorphic net diversification rate = 0.0517, observed 0A monomorphic net diversification rate = 0.0253, Table 2), and a much higher character transition rate from color polymorphism to monomorphism (BiSSE observed 1A color polymorphic transition rate to monomorphic 0A = 0.031, observed monomorphic 0A to 1A color polymorphic = < 0.0001). All null and character-independent SSE models received very little support relative to state-dependent diversification SSE models, accounting for less than 0.1% of the Akaike model weight (Table 1). Complete details of all SSE model net diversification and extinction fraction parameter estimates are given in Table 2. Overall, the HiSSE model accounted for more than 99% of the Akaike model weight and supports our hypothesis that evolutionary transitions to color polymorphism are associated with elevated diversification rates.

### Trait simulations on empirical phylogeny and SSE model adequacy

When we compared AIC values of SSE models on trait data simulated with no correlation to diversification rates on the empirical Lacertidae phylogeny, we found that the BiSSE null model (SSE model with 2 transition rates, equal turnover and extinction fractions, and no hidden states) was selected as the best fit model the most (ΔAIC score = 0, 34.1% of simulations for BiSSE null model). The HiSSE null was the best fit model 21.3% of simulations. Character-independent models, CID-2 and CID-4, were rarely the best fit models on simulated trait data and also had low Akaike model weights compared to other models (0.8% of simulations CID-2 ΔAIC score = 0, 2.9% of simulations CID-4 ΔAIC score = 0, Fig. 3). Character-dependent diversification models, BiSSE and HiSSE, were the best fit models some of the time on simulated color trait data (ΔAIC score = 0, 20.3% of simulations for BiSSE model, 20.6% for HiSSE model, Fig. 3). Ultimately, we recover a Type I error rate (trait-dependent diversification when there should be none) 40.9% of the time when we run the same six SSE models from our observed data on uncorrelated simulated trait data.

### SSE model selection and parameter estimation from posterior distribution of trees

SSE model selection using the empirical color polymorphism data performed on a posterior distribution of 1,000 trees identified BiSSE and HiSSE as the best fit models 96.6% of the time (Fig. 4), which largely supports a color polymorphism trait-dependent diversification scenario in the Lacertidae. The BiSSE and HiSSE models also received larger Akaike model weights than null or character-independent diversification models (Fig. 4). The BiSSE model with no hidden states was selected as the best fit model most often (ΔAIC score = 0, 51.1% of sampled trees), followed by the HiSSE model (ΔAIC score = 0, 45.5% of sampled trees), and the HiSSE null model (ΔAIC score = 0, 4.4% of sampled trees). Of the 44 times the HiSSE null model was selected as the best fit model, it achieved a ΔAIC score > 0-2 28 times. Out of 1,000 potential evolutionary histories of the Lacertidae, the BiSSE null model and character-independent models of diversification were never selected as best fit models.

SSE model net diversification parameter estimates extracted from a subset of 50 phylogenetic trees varied, but overlapped with the estimates obtained from the maximum clade credibility tree (Table 2). Extinction fraction estimates from the posterior subset of trees had higher variance (Table 2).

## DISCUSSION

Color polymorphism research has been dominated by the hypothesis that multiple genetically-based phenotypes can be a precursor to speciation, but with few comparative studies there has been a limited ability to test this hypothesis. We generated a comprehensive family-wide multi-locus species tree of the Lacertidae to elucidate the evolutionary history and macroevolutionary consequences of color polymorphism. Phylogenetic and ancestral state reconstructions of the family Lacertidae suggests the most recent common ancestor of all lacertids was most likely not color polymorphic, and there were probably multiple independent evolutionary transitions to color polymorphism throughout the family tree. We found that the evolution of color polymorphism from monomorphism happens at a much slower rate than evolutionary transitions from color polymorphism to monomorphism. This macroevolutionary-level finding follows empirical results from species-specific case studies that color polymorphism is more easily lost than gained from populations (Corl et al. 2010b; Runemark et al. 2010). Finally, we explored macroevolutionary dynamics within the Lacertidae to test the theory that color polymorphism is a driver of diversification. Amongst several alternative hypotheses that simultaneously consider evolutionary history and trait transitions, we found multiple lines of support for our hypothesis that color polymorphic lineages diversify at a higher rate than monomorphic lineages.

### The Lacertidae: Evolutionary history and color polymorphism

Phylogenies are essential for addressing macroevolutionary hypotheses that use interspecific data, and a long-contested phylogeny has limited our understanding of the evolutionary history and macroevolutionary dynamics of the Lacertidae (Fu 2000). Molecular investigations of the entire family Lacertidae at the species level are rare, and topological relationships in the family tree remain controversial (Fu 1998; Harris et al. 1998; Fu 2000; Arnold et al. 2007; Mayer & Pavlicev 2007; Garcia-Porta et al. 2019). Phylogenetic uncertainty in the Lacertidae likely stems from a combination of early and recent bursts of diversification (Fu 2000; Garcia-Porta et al. 2019). Family-level phylogenies of lacertids usually recover low support deep within the tree and at nodes connecting short branches near the tips (Fu 2000; Arnold et al. 2007; Garcia-Porta et al. 2019). We find a similar pattern in our data, with an early period of rapid diversification deep within the tree and low support or non-traditional placement of short branch taxa. Fossils and genetic data from Europe help resolve relationships among morphologically convergent European lacertids, but limited genetic data and poor fossil records in other areas where lacertids currently occur, such as the Middle East, Africa, and Asia, hinder our ability to generate phylogenies with strong support deep in the tree and between subgroups in these lineages (Hipsley et al. 2009). In particular, Eurasian lacertids and species with expansive geographic distributions tend to have uncertain placement in phylogenetic reconstructions (Fu 2000; Garcia-Porta et al. 2019). Twenty years later, and with far more molecular markers, we echo Fu’s (2000) sentiment that to resolve nodes with low support in the lacertid family tree, future investigations should focus on interrogating species-level evolutionary relationships between a few widely distributed and contested genera that are probably not monophyletic (e.g.: the highly polyphyletic former Lacerta genus; Arnold et al. 2007).

Accommodating phylogenetic uncertainty is essential to evolutionary studies. Indeed, the one true evolutionary history escapes us due to missing data from past processes such as extinction and an incomplete fossil record, and from limitations in the present from missing data, taxonomic uncertainty, and the continuous and ephemeral nature of speciation (Huelsenbeck et al. 2000; Rosenblum et al. 2012; Louca & Pennell 2020). We account for phylogenetic disagreement and uncertainty with trait simulation studies, test suites of alternative hypotheses on 1,000 sampled trees from our Bayesian posterior, and run parallel comparative methods analyses on our own lacertid tree inference and that of others (Garcia-Porta et al. 2019). Through these methods, we find multiple lines of evidence all in agreement with regard to the evolutionary history of color polymorphism in the Lacertidae. Whether the Gallotinae truly belong sister to all Lacertinae (Arnold et al. 2007; Garcia-Porta 2019), or somewhere nested within (Fu 2000), the phylogenetic structure of the evolution of color polymorphism across extant lacertids places the first instances of color polymorphism deep within the tree, though most likely not at the ancestor of all lacertids (Fig. 2).

The Lacertidae exhibit a high degree of color polymorphism, spanning 10 genera and comprising 43 species that share a similar throat color polymorphism. Phylogenetic and ancestral state reconstruction analyses revealed that the ancestor of all lacertids was probably not color polymorphic, and that color polymorphism has been gained and lost several times throughout the evolutionary history of the Lacertidae. That the ancestor of all lacertids was most likely not color polymorphic is not surprising, given that the closest relatives of lacertids used as outgroup taxa are themselves not color polymorphic. Color polymorphism, however, appears to have evolved relatively early in the history of lacertids, during or shortly after an initial early period of diversification in the family. Color polymorphism seems to be a trait restricted to the Lacertini tribe, and most likely evolved in the group. Our results that recover an ancient origin of color polymorphism in the group also underline recent findings from a study of the highly polymorphic *Podarcis* group, which found patterns of molecular evolution at the color polymorphism pigmentation loci that indicate the alleles are of ancient origin (Andrade et al. 2019). Contrary to our predictions, we estimate that color polymorphism evolved more than once throughout the evolutionary history of the Lacertidae, possibly up to seven times. This result is surprising given that empirical studies of color polymorphic taxa at the species level report that morph loss in populations represents lost genetic variation that cannot likely be regained (Corl et al. 2010b). Though, our estimates indicate the rate of loss far exceeds the rate of gain of color polymorphism, which aligns with our expectations. The tendency for color polymorphism to be lost faster than it evolves follows from theory on morphic speciation (West-Eberhard 1986; Gray & McKinnon 2007). If color polymorphic species have highly variable or large geographic ranges where gene flow between populations is infrequent, populations that experience morph loss may diverge quickly genetically and phenotypically (Corl et al. 2010a,b), setting the stage for speciation. This scenario would generate a phylogenetic pattern where color polymorphic lineages give rise to daughter lineages that are monomorphic, which we see in our ancestral state reconstruction in the *Apathya, Darevskia, Dinarolacerta, Iberolacerta*, and *Phoenicolacerta* generic groups (Fig. 2).

An unexpected pattern emerges in the speciose *Podarcis* clade, where 19 out of 23 extant species are color polymorphic. If color morph loss or divergence progresses to speciation, we would expect to see polymorphic lineages give rise to monomorphic descendant lineages (Jamie & Meier 2020). Further, the only other comparative study on color polymorphism and diversification we are aware of found that color polymorphic lineages tend to be younger than monomorphic lineages (Hugall & Stuart-Fox 2012), which also aligns with early theoretical models of morphic speciation driven by morph loss and fixation (West-Eberhard 1986). However, the *Podarcis* group is not particularly young compared to other groups in the Lacertidae, nor does it exhibit many short branches like the color polymorphic *Darevskia* clade. So what might explain persistent polymorphism in *Podarcis*? A recent genomic study that identified genes controlling color differences amongst morphs in *Podarcis muralis* also found some evidence for inter-specific color allele sharing with other *Podarcis* species (Andrade et al. 2019). To retain color polymorphism after speciation, the alleles for different morphs must be present, either ancestrally or arising again through a novel mutation. Ancestral genetic variation may persist past speciation via introgression or standing ancestral variation (Andrade et al. 2019; Jamie & Meier 2020). If speciation rates are high, there may be more opportunity for introgressed morph alleles and thus morph persistence past speciation if the ancestral color polymorphic lineage comes into secondary contact with a monomorphic lineage (Jamie & Meier 2020). Taken together, these genomic and phylogenetic investigations of the *Podarcis* clade raise interesting implications for the role of hybridization and introgression in the evolution and long-term maintenance of color polymorphism and its relationship to speciation rates (Jamie & Meier 2020).

### Color polymorphism is associated with rapid diversification rates

Theory suggests that dramatic intra-specific phenotypic diversity and the underlying processes that maintain it may promote rapid speciation in color polymorphic lineages (West-Eberhard 1986; Gray & McKinnon 2007; Forsman et al. 2008). We find evidence for this in the Lacertidae, where diversification rates are substantially faster in lineages in the color polymorphic state. Across many possible phylogenetic reconstructions of the Lacertidae, we consistently estimate that net diversification is almost double the rate in color polymorphic lineages than monomorphic lineages. Indeed, lacertids also exhibit faster character transition rates from color polymorphism to monomorphism, which is consistent with theory that suggests speciation occurs by morph loss and fixation of remaining morphs (West-Eberhard 1986). Trait simulations and state-speciation extinction models run on many possible trees, including a tree inferred with a large phylogenomic dataset, suggest that it is unlikely that the shape of the Lacertidae phylogeny produces false estimates of trait-dependent diversification. These findings are also supported by empirical studies that show repeated loss, fixation, and rapid divergence of morph types among populations of color polymorphic species (Corl et al. 2010b).

Animal color and pattern are important traits involved in processes such as mate choice, species recognition, and sexual selection, which can all play a role in accelerating speciation (Houde & Endler 1990; Roulin 2004). Across many color polymorphic species, morph color is often involved in intra-specific visual signaling to communicate myriad messages in a variety of social and environmental contexts (Gray & McKinnon 2007). In social contexts, morph color can indicate reproductive strategy (Sinervo & Lively 1996) and morph color is often a factor in mate choice (Pryke & Griffith 2007). In birds, reptiles, and fish, the prevailing environment and lighting conditions affect the efficacy and transmission of signals displayed by different color morphs (Gray & McKinnon 2007), and color morphs may segregate microhabitat to optimize signal transmission (Endler 1984). Thus, if sexual and/or natural selection pressures shift away from balancing color polymorphism toward favoring the phenotype of one or several morphs over another, divergent or directional selection could result in morph loss from a population. Because color polymorphic species inherently possess extreme variation, there exists increased opportunity for selection or drift to operate against any one of several distinct color morph phenotypes, which may explain elevated diversification rates in color polymorphic lineages. Theoretical expectations and empirical studies of populations show that morph loss and fixation can both result in rapid divergence (West-Eberhard 1986; Corl et al 2010a,b), but the microevolutionary processes operating within and between populations that disrupt balanced color polymorphisms and generate divergence remain less understood and generalizable. Further study is needed to quantify the relative roles of natural selection, sexual selection, and drift in color polymorphism maintenance and speciation.

Ultimately, the color polymorphic condition in lacertids is associated with elevated diversification rates. Species have many traits, and it is unlikely that a single trait is the only factor that accounts for increased or decreased diversification rates. Indeed, the best fit model was trait-dependent diversification that included additional unobserved “hidden states” that are correlated with color polymorphism. Color polymorphism is usually accompanied by alternative morph-specific ecological, morphological, physiological, and behavioral syndromes, or correlated traits (Lattanzio & Miles 2016; Huyghe et al. 2009a,b; Sinervo & Lively 1996; Sinervo & Svensson 2002). Here, color and other heritable traits are likely subject to multivariate selection, wherein correlational selection builds up genetic correlations through linkage disequilibrium at loci underlying the traits (Sinervo & Svensson 2002). Correlation between color morphs and traits related to fitness, such as reproductive strategy (Sinervo & Lively 1996; Galeotti et al. 2013), or reproductive hormone levels (Huyghe et al. 2009b), or body size, can produce color morphs with alternative adaptations that occupy different adaptive peaks (West-Eberhard 1986). When a color polymorphic lineage with multiple balanced adaptive peaks faces strong or novel selective forces, one or more of the peaks may shift, and a morph or morphs must cross “valleys” of selection to persist else they are lost. Thus, the nature of color polymorphism and correlated traits could increase the potential for color polymorphism to contribute to divergence amongst morphs and speciation.

### Color polymorphism: linking micro-evolutionary process and macroevolutionary pattern

Color polymorphism is widespread throughout the tree of life, but our understanding of the mechanisms underlying morph evolution and the processes that influence the shape of the tree remain fragmented. The evolutionary mechanisms that maintain alternative color phenotypes within a population are also likely involved in morph loss, divergence, and speciation (Gray & McKinnon 2007). These population-level processes, particularly the effects of natural and sexual selection and their relative roles within and between populations, are of utmost interest (Jamie & Meier 2020). Color polymorphism is often studied at the level of a single species (Huyghe et al. 2009a,b; Corl et al. 2010a,b; Runemark et al. 2010), but macroevolutionary perspectives will provide deeper insights into the origin, duration of maintenance, and inter-specific persistence of color polymorphism (Gray & McKinnon 2007; Jamie & Meier 2020). Very few empirical studies link population-level evolutionary processes and the evolution of color polymorphism at the macroevolutionary scale (Hugall & Stuart-Fox 2012; Willink et al. 2019). Studies that investigate trait variation within species and use biologically meaningful versions of those inputs at the between-species level with the use of comparative methods are needed (Gray & McKinnon 2007). Such investigations will illuminate the connection between evolutionary process and macroevolutionary pattern. Color polymorphisms offer ideal model systems to study how micro-evolutionary dynamics shape macroevolutionary patterns of diversification. Our results show that the Lacertidae, in particular, offer a promising avenue for interrogating the relative contributions of different forms of selection on alternative phenotypes, and how population-level color morph dynamics scale up to influence macroevolutionary patterns.

## ACKNOWLEDGEMENTS

We would like to thank Jessica Blois and Jeremy Beaulieu for their helpful comments during the preparation of our manuscript and Indiana Madden for assistance in compiling phenotypic data for this project. Computation for phylogenetic analyses was made possible with assistance from Sarvani Chadalapaka and the many cores of the MERCED cluster (National Science Foundation Grant No. ACI-1429783). We would also like to thank the Herpetology Collections at the Zoologisches Forschungmuseum Alexander Koenig (Morris Flecks) and the University of Michigan Museum of Zoology (Gregory Schneider) for access to lacertid specimens. This work was supported by a Graduate Student Research Award from the Society of Systematic Biologists awarded to K.M. Brock.

